# Using Maternally Inherited Haploid Tissue to Resolve Parental Alleles: Investigating Genomic Imprinting in Scots Pine (*Pinus sylvestris*)

**DOI:** 10.64898/2026.03.24.713999

**Authors:** Robert Kesälahti, Sandra Cervantes, Alina K. Niskanen, Tanja Pyhäjärvi

## Abstract

Genomic imprinting is a rare epigenetic phenomenon in plants and animals, defined by parent-of-origin specific gene expression. Its molecular mechanisms and evolutionary significance remain incompletely understood. In this study, we investigated whether genomic imprinting occurs in Scots pine and, by extension, in other conifers to gain insight into the evolutionary origins of imprinting. We performed reciprocal crosses to assess imprinting in seed embryos and applied a unique approach that used exome-capture data from the haploid, maternally inherited megagametophyte tissue to identify maternal alleles, thereby allowing us to infer paternal alleles in the embryos of the same seeds. Our findings show that maternally inherited haploid megagametophyte tissue offers an effective strategy for resolving parental alleles in offspring while simultaneously removing extensive paralogous variation from the dataset. This framework is broadly applicable to other conifer species and to taxa that possess comparable maternally derived haploid tissues. No evidence of genomic imprinting was detected. Although the limited overlap between the exome-capture and RNA-sequencing datasets and the stringent paralog filtering reduced the amount of analyzable data considerably, the absence of detectable imprinting may also reflect genuinely weak or absent imprinting signals in conifers. We identified several limitations in this preliminary study and outline recommendations for future work to overcome them, and additional research will be necessary to determine whether genomic imprinting occurs in conifers

## 1 Introduction

Genomic imprinting, a rare epigenetic phenomenon observed in both plants and animals and affecting only a small subset of genes, is characterized by the parent-of-origin–specific expression of genes. In practice, this means that an allele may be silenced when paternally inherited, resulting in monoallelic or near monoallelic expression, or conversely, when maternally inherited. Imprinting arises through epigenetic modifications, including DNA methylation (Bourc’his et al., 2001 and histone marks (Ciccone et al., 2009), and is further regulated by the roles of noncoding RNAs (Sleutels et al., 2002) and RNA interference mechanisms (Vu et al., 2013). These epigenetic marks are not observable through DNA sequence analysis alone. This phenomenon is highly fascinating, as it bends the rules of Mendelian inheritance by exerting hidden control over allele-specific gene expression and providing a means to temporarily bypass potential disadvantages of diploidy. Although imprinting has been recognized for decades, its underlying mechanisms and evolutionary significance remain incompletely understood, leaving many questions unresolved.

In 1970, J.L. Kermicle discovered the first imprinted locus, noting that R-mottling, a pattern of uneven anthocyanin distribution, was only present in the aleurone layer of maize endosperm when the pollen parent had an RR genotype, but absent when the rr genotype was the pollen parent in RR x rr crosses (Kermicle, 1970). Kermicle concluded that dosage effects could not explain these observations. Subsequent studies identified the first signs of imprinting in mice (Johnson, 1974) and in humans (Lubinsky et al., 1974), laying the groundwork for later theories about its evolutionary origin. The first clear evidence that imprinting affects seed development came in 1999 with the discovery of the MEDEA gene, which encodes a Polycomb group (PcG) protein and is expressed only from the maternal allele in *Arabidopsis thaliana* endosperm (Vielle-Calzada et al., 1999). Mutations in the maternal copy cause seed abortion and abnormal endosperm and embryo development. This finding showed that genomic imprinting is essential for normal seed development. Since then, extensive genome-wide imprinting studies have been carried out in various angiosperm species, including *A. thaliana* (Gehring et al., 2011; Wolff et al., 2011), rice (Luo et al., 2011), maize (Waters et al., 2013), castor bean (Xu et al., 2014), and Brassicaceae (Hatorangan et al., 2016), revealing hundreds of imprinted loci and highlighting the prevalence of this phenomenon.

However, research into genomic imprinting has predominantly focused on angiosperm species within the plant kingdom. This is because most imprinted genes have been discovered in the triploid endosperm, a characteristic unique to angiosperms. The widely accepted Kinship theory proposes that imprinting originates from paternal conflict over resource distribution for the developing offspring and may have evolved independently in both angiosperms and animals through convergent evolution (Feil and Berger, 2007; Haig, 2000; Haig and Westoby, 1989). Early studies from angiosperms and mammals have supported this hypothesis (Leighton et al., 1995; Scott et al., 1998). However, other theories, such as sexual conflict (Day and Bonduriansky, 2004) and maternal-offspring co-adaptation (Wolf and Hager, 2006) have been proposed due to inconsistencies with the kinship theory. Several observations contradict the kinship theory. First, not all imprinted genes are involved in growth or nutrient transfer (Gehring et al., 2011; Pignatta et al., 2014). Second, imprinting is not exclusive to the endosperm, as it has also been observed in embryos (Jahnke and Scholten, 2009; Raissig et al., 2013; Waters et al., 2011). Third, there are different types of imprinting, some of which have been recognized for a long time. For instance, in certain insects such as sciarids, male sex chromosomes or even the entire paternal haploid set are imprinted as part of the sex determination process (reviewed in Sánchez, 2014), a phenomenon known as paternal genome elimination that was first described by Du Bois in 1933 (Du Bois, 1933). A recent discovery in the non-vascular plant *Marchantia polymorpha*, which belongs to the liverwort group, reveals that the entire paternal genome is repressed during the diploid phase of its life cycle (Montgomery and Berger, 2024). This extensive paternal genome imprinting results in embryos being functionally haploid and under strong maternal control. Genomic imprinting was also recently discovered in hybrids of two water lily sister species (Povilus et al., 2025). Water lilies and other Nymphaeales predate the monocot–eudicot divergence and are distinguished from other angiosperms by their unique diploid endosperm and maternal storage tissue (perisperm). Findings from Povilus et al., (2025) suggest that genomic imprinting was already present at the origin of endosperm and, together with evidence from *Marchantia polymorpha*, point to the possibility that imprinting is more widespread across the plant kingdom than previously recognized. To fully comprehend the evolutionary origins and selective pressures driving genomic imprinting, it is necessary to conduct studies across various plant lineages.

Despite the theoretical framework and empirical evidence suggesting that genomic imprinting could be more prevalent, there remains a significant gap in our understanding of this phenomenon in non-angiosperm plant species. Conifers, one of the four extant lineages of Gymnospermae, are characterized by exceptionally long generation times and repeat-rich genomes that rank among the largest in the plant kingdom (De La Torre et al., 2014). Their slow rates of genome turnover create a genomic environment that contrasts sharply with that of angiosperms, with implications for the evolution of epigenetic marks, gene silencing, and regulatory divergence. In conifers, the embryo develops for an extended period within a long-lived haploid female gametophyte, creating a distinctly different context for potential parental conflict, resource allocation, and epigenetic regulation. Conifers also exhibit strong local adaptation, reflected in their dominance across vast forest biomes (Savolainen et al., 2007). Maternal environmental effects have been proposed as one mechanism through which mothers can transmit locally adaptive cues to their offspring (Galloway, 2005). This could be especially relevant in wind-pollinated species, such as conifers, where seeds typically disperse only short distances and offspring often encounter environments similar to those of the maternal plant (Vander Wall, 2023). Genomic imprinting has been suggested as one potential mechanism contributing to such parent-of-origin specific influences on offspring phenotype. As early-diverging seed plants, conifers occupy an intermediate position between early angiosperms, such as water lilies, and earlier land-plant lineages, such as liverworts, making them a crucial missing piece for understanding how genomic imprinting evolved across the plant kingdom.

Traditional studies of genomic imprinting have involved reciprocal crosses between two highly homozygous inbred lines, resulting in offspring with a high level of heterozygosity. This heterozygosity is crucial for distinguishing between parental alleles and assessing their imprinting status. The use of inbred lines simplifies the tracking of allele origins. Nonetheless, highly inbred lines are difficult, if not impossible, to generate in naturally outcrossing species or in species with long generation time. Another significant limitation for non-model species has been the lack of high-quality reference genomes, which is critical for imprinting studies where even minor biases can lead to incorrect interpretations. The recent publications of high-quality reference genomes for species such as *Pinus tabuliformis* (Niu et al., 2022), *Pinus densiflora* (Jang et al., 2024), *Pinus sylvestris* (Nilsson et al., 2025), and *Picea abies* (Nilsson et al., 2025) have opened new avenues for research and analysis in conifers. However, a major technical challenge in these species is the abundance of paralogous regions, arising from large multicopy gene families, duplicated genes, and numerous pseudogenes. Paralogs pose substantial problems to mapping and genotyping even in highly selfing species such as *A. thaliana* (Jaegle et al., 2023). Consequently, studying imprinting in conifers requires carefully balancing stringent filtering to remove paralogs against the risk of discarding genuinely imprinted sites

In this study, we conducted, to our knowledge, the first genomic imprinting analysis on conifers, examining Scots pine (*Pinus sylvestris*) embryos obtained from reciprocal crosses between tree pairs from a wild population. We utilized a novel approach to study imprinting by using exome-capture data from the haploid, maternally inherited megagametophyte to identify maternal alleles and infer paternal alleles in the corresponding embryo. We also used the haploid tissue to identify and remove paralous sites from data. This allowed us to investigate how imprinting may contribute to adaptive gene-expression regulation in Scots pine’s large, repeat-rich genome

## 2 Materials and methods

### 2.1 Reciprocal crosses and seed preparation

Six *Pinus sylvestris* trees (251, 320, 397, 443, 463 and 485) from a Finnish Punkaharju ISS population (ISS, https://www.evoltree.eu/resources/intensive-study-sites/sites/site/punkaharju) were chosen for re-ciprocal crosses. The Natural Resources Institute Finland (LUKE) executed reciprocal crosses among pairs: 251×320, 397×443, and 463×485. Five seeds were randomly selected from each direction of a reciprocal cross, yielding 10 seeds per cross and 30 seeds in total for the experiment. The seeds were placed in a controlled environment room (23°C, 70% RH, continuous light at ≈ 500 *µ*m *m*^2^*/s*^2^) to germinate for 48 hours on small plastic Petri dishes lined with four layers of filter paper saturated with distilled water. Subsequently, the seeds were then dissected under a microscope to separate embryonic tissue from megagametophytic tissue.

### 2.2 RNA-sequencing of embryos

Messenger RNA was isolated from the embryonic tissue utilizing Dynabeads mRNA DIRECT Micro Kit (Thermo Fisher Scientific) as directed by the manufacture’s instructions. The RNA samples were then quantified with the RNA 6000 Pico Kit (Agilent) and Bioanalyzer 2100 (Agilent). Subsequently, the RNA libraries were constructed using NEBNext Ultra Directional RNA Library Prep Kit for Illumina (New England Biolabs) together with the NEBNext Multiplex Oligos for Illumina (Index Primers Set 1, New England Biolabs). The size distribution of the DNA fragments in the libraries was determined using High Sensitivity DNA Kit (Agilent) and Bioanalyzer 2100. Quantification of the libraries was performed with NEBNext Library Quant Kit for Illumina (NEW ENGLAND Biolabs) and LightCycler 480 (Roche), followed by sequencing on NextSeq500 using a Mid Output kit at the Biocenter Oulu Sequencing Center. Out of the 30 samples, one failed during sequencing, resulting in a total of 29 samples being sequenced.

### 2.3 Exome capture from haploid megagametophyte tissue

DNA was extracted from the megagametophytic tissue using E.Z.N.A SP Plant DNA Kit (Omega Bio-tek) according to the manufacturer’s manual. An additional sodium hydroxide treatment was applied to the HiBind DNA Midi Column prior to sample loading and a double-elution process was performed to enhance DNA yield. DNA concentration was quantified using Quant-iT PicoGreen dsDNA Kit (Thermo Fisher Scientific). Samples with DNA concentrations exceeding 12 ng/*µ*l were diluted to 12 ng/*µ*l using TE buffer (1 mM Tris–HCl, pH 8.0, 0.01 mM EDTA) to control fragment size during fragmentation. DNA was fragmented using a Bioruptor UCD-200 (Diagenode) for 3 × 10 min plus an additional 5 min in 30 s on/30 s off cycles at power setting ‘L’. Samples were vortexed between each round of fragmentation. Size selection for the fragments was performed using Agencourt AMPure XP beads (Beckman Coulter) to obtain fragments within the 250–450 bp range. A bead-to-sample volume ratio of 0.6 was used for left-side selection and 0.133 for right-side selection. Concentration of the fragmented DNA was re-measured using the Quant-iT PicoGreen dsDNA Kit (Thermo Fisher Scientific) to estimate DNA loss during the size-selection process, and the size distributions were quantified with the High Sensitivity DNA Kit (Agilent) on a Bioanalyzer 2100. Libraries were constructed using KAPA HyperPrep Kit (Roche) with SeqCap Adapter Kit A and B (Roche) indices. Due to low DNA concentrations in the samples, only half of the recommended volumes for the end repair, A-tailing, and adapter ligation steps were used. Two multiplexed pools containing 1000 ng of library DNA were prepared. Exome captures for the two multiplexed pools were performed using a SeqCap EZ HyperCap (Roche) kit with a custom bait set: SeqCap Design PiSy UOULU (Kesälahti et al., 2025). Species-specific c0t-1 DNA (30,000 ng) was used in the hybridization reaction to improve capture of unique exons and reduce capture of tandem repeats. C0t-1 DNA was prepared as described in Kesälahti et al., 2025. The size distribution of the captured libraries was analyzed using High Sensitivity DNA Kit with Bionanalyzer 2100 and quantified using Kapa Library Quant Kit (Roche) with LightCycler 480 and sequenced by Novogene. Two samples failed sequencing due to issues with the indices, resulting in a total of 27 sequenced samples.

### 2.4 Bioinformatic workflow

RNA-sequencing data were utilized to identify genes exhibiting allele-specific expression in embryos. Exome capture data from the maternally inherited megagametophytes, whose haplotype within a seed is always identical to the maternal haplotype of the embryo, were used to identify maternal alleles of genes exhibiting allele-specific expression. Analysis included 27 sample pairs, comprising 9 samples from each of the reciprocal crosses. The scripts used for the bioinformatic analyses described below are available at (link)

Preprocessing and quality control of all FastQ files was performed using fastp v0.24.0 (S. Chen, 2023). Fastp reports were aggregated using MultiQC v1.9 (Ewels et al., 2016).

#### 2.4.1 Identification of heterozygous SNPs and extraction of read counts

RNA-sequencing reads were mapped to the reference genome of *P. sylvestris* (Nilsson et al., 2025) using STAR v2.7.11b (Dobin et al., 2012). Duplicate reads were removed with Picard MarkDuplicates and read group tags were added using Picard AddOrReplaceReadGroups tool (Picard Tools) v2.27.4; Broad Institute, 2022). Unique alignments were retained by filtering out multi-mapped reads based on the NH:i tag. Alignments with NH:i values greater than one were removed using Samtools version 1.9 (Danecek et al., 2021) view command in combination with grep. Filtered BAM files were indexed using SAMtools index command. CSI indices were selected for indexing due to their ability to manage chromosomes larger than 512 Mbp. A joint variant call was performed across all samples to identify heterozygous single nucleotide polymorphisms (SNPs) using BCFtools version 1.9 (Danecek et al., 2021) mpileup and call commands. The ASEReadCounter tool from Genome Analysis Tool Kit (GATK) version 4.5.0.0; (McKenna et al., 2010) was used to count reads for the reference and the alternative alleles at each heterozygous SNP site. Prior to counting, indels were filtered out and an indexed VCF file was created. Indexing of the reference genome was completed using Samtools faidx command, and a sequence dictionary was generated with GATK’s CreateSequenceDictionary tool. Indels were removed using BCFtools view command. Sites that did not show any reads from one of the two alleles were excluded using the Unix awk command. These sites were present because they had been detected as heterozygous in at least one other sample during the variant joining process, but the sample in question was not heterozygous for that particular site.

#### 2.4.2 Identification of maternal alleles

Exome capture reads were mapped to the reference genome of *P. sylvestris* using the bwa-mem algorithm (bwa v.0.7.17;(Li, 2013) with the default parameters. Duplicate reads were removed with Picard MarkDuplicates tool and then indexed using SAMtools index command. Separate haploid variant calling was performed for each BAM file, utilizing BCFtools mpileup and call commands, and indels and VCF headers were removed using BCFtools view and filter commands. A substantial number of heterozygous sites were detected in the haploid variant calls, likely caused by reads from paralogous genes mapping to the same genomic regions. To exclude these positions, a minimum depth of two was imposed for filtering using BCFtools filter command. Heterozygosity was determined from all haploid variant calls using the DP4 field, considering a site heterozygous if it had at least one read for the reference allele and one for the alternative allele. All heterozygous sites from variant call outputs were removed regardless of their presence in a sample, to account for potential paralogous read origin. Maternal alleles for heterozygous SNPs from embryo RNA-sequencing data were identified by merging the filtered haploid variant call tables with ASEReadCounter output tables based on chromosome and position, using the Unix join command.

#### 2.4.3 Genotype-based filtering

A matrix of maternal genotypes was constructed for each heterozygous SNP identified in a reciprocal cross. The SNPs were then subjected to a two-step filtration process. First, the offspring genotypes had to be the same across all samples within a cross, meaning that all samples in the AxB cross had to share the same genotype, and all samples in the BxA cross had to share the same genotype, which was distinct from the AxB cross. Second, we required each site to have genotype data in the majority of samples within a cross—at least five of the nine individuals—to exclude positions with extensive missing data. By focusing on heterozygous SNPs that displayed a consistent genotype across all samples in each cross, the imprinting analysis was narrowed down to sites that were genuinely heterozygous. This conservative filtering approach was used to reduce uncertainty in assigning parental alleles, as no reliable method exists to distinguish true heterozygous sites from those arising from paralogous mapping.

#### 2.4.4 Genomic imprinting analysis

Genomic imprinting was examined using the statistical approach introduced by Wyder et al. (2019). This methodology uses generalized linear models (GLM) and a negative binomial distribution implemented in edgeR (Y. Chen et al., 2025). For a full description of the method, see (Wyder et al., 2019). The analyses were conducted in R version 4.5.0 using edgeR version 4.6.3, using a modified script from Wyder et al. (2019). Data visualizations were generated using both the ggplot2 package (Wickham, 2016) and base R graphics (R Core Team, 2025). Each reciprocal cross was analyzed independently, incorporating all biological replicates within a cross. A heterozygous SNP was included in the analysis only if at least two samples per cross had a minimum of ten reads from the maternal or paternal allele. This step excludes SNPs with very low expression while retaining the ability to detect genomic imprinting with both mono-allelic and biased expression patterns. The top ten heterozygous SNPs associated with genomic imprinting were ranked by their raw P-values from each reciprocal cross. These SNPs were then annotated against the *P. sylvestris* reference genome functional annotation file using BEDtools v.2.31.1 (Quinlan and Hall, 2010) with the intersect command. Biological functions were inferred by matching the mRNA level annotations of heterozygous SNPs to the longest representative annotation for each mRNA sequence (Nilsson et al., 2025). The mRNA-level annotations were retrieved in FASTA format using Bedtools getfasta command.

## 3 Results

To identify genes exhibiting genomic imprinting, we analyzed reciprocal crosses among three Scots pine tree pairs from a wild Finnish population. From these crosses, short-read RNA-sequencing data (2×150 bp) were obtained from 27 embryos, and exome capture data (2×150 bp) from 27 haploid megagametophytes of the same seeds. A custom exome capture bait set (SeqCap Design PiSy UOULU) targeted 18,516 regions, spanning a total capture space of 9,345,488 bp. Both types of sequencing libraries showed substantial variation in the number of raw of reads, which was reflected in the number of mapped reads and PCR duplicates (Supplementary Table 1). From the RNA-sequencing data, we identified heterozygous SNPs (het-SNPs) and extracted their allele-specific read counts. Across samples, the number of het-SNPs ranged from 150,627 to 370,528 (Table 1). We used the haploid megagametophyte data from exome captures to remove paralogous sites and identify parental alleles. All-site haploid variant call was performed for each megagametophyte sample separately and sites containing reads for two different alleles were removed. Using this approach we were able to remove 40,351,260 paralogous sites from our data. After the paralog removal, we had between 48,787,750 and 82,742,737 sites available for maternal allele identification per haploid sample (Supplementary Table 1). Maternal alleles were identified by finding chromosomal positions where heterozygous SNPs from RNA-sequencing data and haploid all-sites variant call data overlapped. This method allowed for the identification of maternal alleles for between 1.3 and 3.6 percent of heterozygous SNPs (Table 1), indicating low overlap between the two different datasets. Further paralog filtering was applied, retaining only heterozygous SNPs with identical maternal alleles among biological replicates within each reciprocal cross. Additionally, maternal allele data was required from the majority of samples (5/9 for each reciprocal cross) to ensure reliable genomic imprinting analysis. After filtering, between 348 and 845 sites per sample were available for imprinting analysis (Table 1).

**Table 1:**
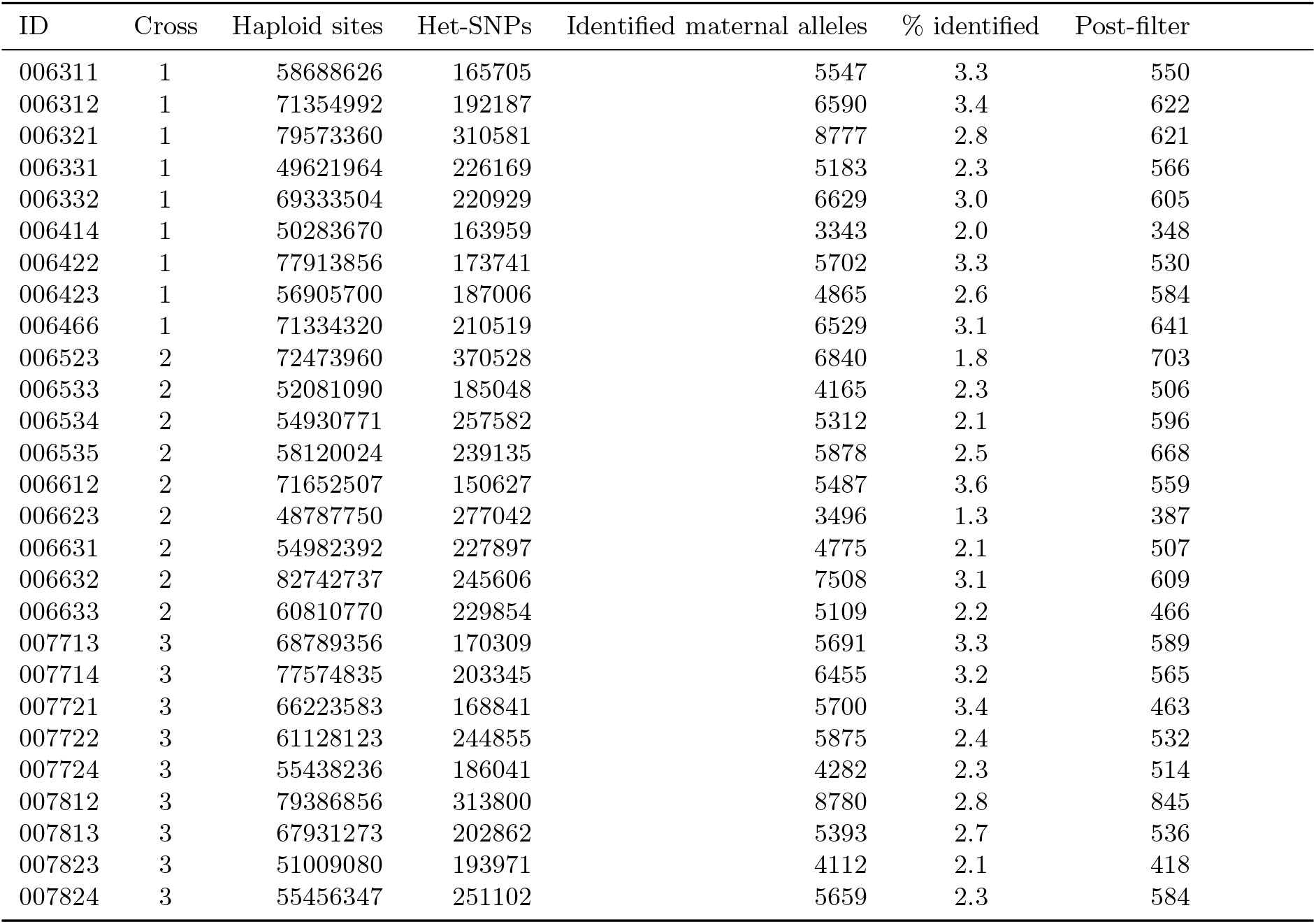
Summary of maternal allele identification and genotype filtering results.

We implemented an edgeR-based pipeline, which employs generalized linear models (GLM) and a negative binomial distribution to assess genomic imprinting from our het-SNPs. This method has been demonstrated to outperform various other techniques (Wyder et al., 2019). The imprinting analysis was conducted separately for each of the three reciprocal crosses, including all biological replicates within each cross. Across all biological replicates, a set of 1317, 1297 and 1244 unique het-SNPs were available for the analysis. To exclude het-SNPs with very low expression levels, a filtering step requiring that at least two samples per cross had a minimum of ten reads from either the maternal or paternal allele was incorporated. After filtering, 139, 184, and 184 heterozygous SNPs remained for imprinting analysis in the three reciprocal crosses. The limited numbers highlight that the majority of het-SNPs were characterized by very low expression levels. Additionally, substantial variation in expression levels was observed among biological replicates. All three reciprocal crosses show a similar increase in biological coefficient of variation (BCV) values with higher expression levels (see Figure 1). BCV measures the inherent biological variability in gene expression among replicates while accounting for technical variation. The common BCV values, close to 1.5, indicate high biological variability across reciprocal crosses. The common dispersion values for each cross (2.30, 2.17, and 1.70) are also high, showing that the dataset has substantial variability.

**Figure 1.**
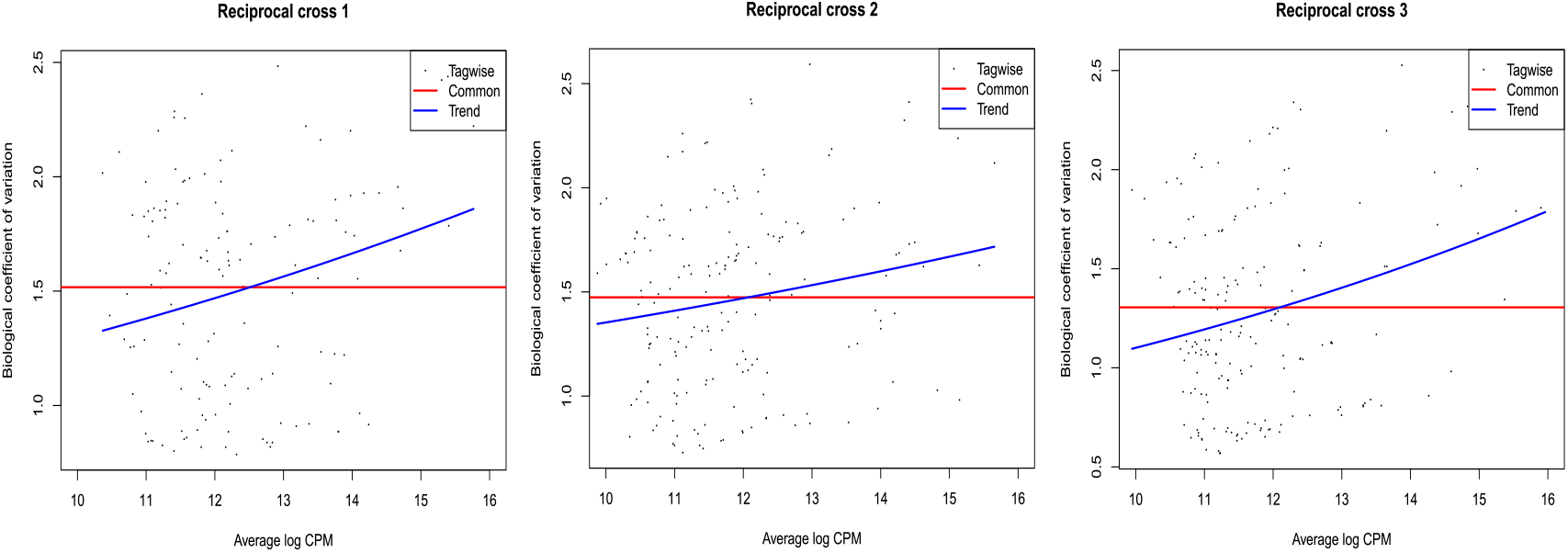
Biological coefficient of variation (BCV) plots from edgeR for three reciprocal crosses. BCV plots show the relationship between the square root of the biological coefficient of variation (y-axis) and the average log-counts per million (log-CPM) for each heterozygous SNP (x-axis). Each plot includes the estimated common BCV (red line), which provides a global dispersion measure used in downstream genomic imprinting analyses. The blue line represents the trended dispersion estimate, capturing the mean–variance relationship across het-SNPs.

Although expression levels varied considerably among replicates, the dataset still provided a homogeneous representation of the Scots pine genome, as the het-SNPs were evenly distributed across all twelve chromosomes (Figure 2), offering a genome-wide basis for assessing parental expression bias. Parental bias is fairly balanced in both directions and shows no major differences among chromosomes (Figure 2), with most logFC values being small. Despite a handful of het-SNPs showing stronger allelic biases and appearing as potential imprinting candidates, the analysis did not identify any statistically significant maternal or paternal bias in any of the reciprocal crosses after the FDR correction (Table 2). Two het-SNPs in the second reciprocal cross showed raw P-values below the 0.05 threshold, but neither passed the false discovery rate (FDR) cutoff of 5%. The analysis produced inflated FDR values across all het-SNPs, indicating absence of convincing evidence of parental biased expression. Most raw P-values were high (0.75–1.0), which in turn contributed to the inflated adjusted FDR values (Figure 3). Closer inspection of the two het-SNPs with significant P-values revealed that they are located in the same mRNA (PS chr01 G002762.mRNA.1) associated with plant stress response, particularly to drought (González and Iusem, 2014; Jing et al., 2017). However, in closer inspection the expression patterns of this mRNA do not consistently align with imprinting: it shows high mono-allelic expression from the maternal allele in one biological replicate in the first cross, slight paternal bias in other replicates, and in the second cross, consistently higher maternal expression across all replicates (Figure 4). Interestingly, a het-SNP associated with an RNA recognition motif (RRM) containing gene (PS chr11 G046551.mRNA.1) shows an expression pattern in the second cross that resembles genomic imprinting, where the maternal allele consistently has higher expression. The P-value for this SNP (0.077) is slightly above the significance threshold, likely because expression data are available from only three replicates in the first cross and two in the second. RRM-containing proteins are abundant and widespread in eukaryotes and are involved in plant development and stress responses (Shi et al., 2017). They form large gene families in plants and display a high degree of sequence conservation within their encoded domains, as recently demonstrated in rice (Jiang et al., 2024).

**Table 2:**
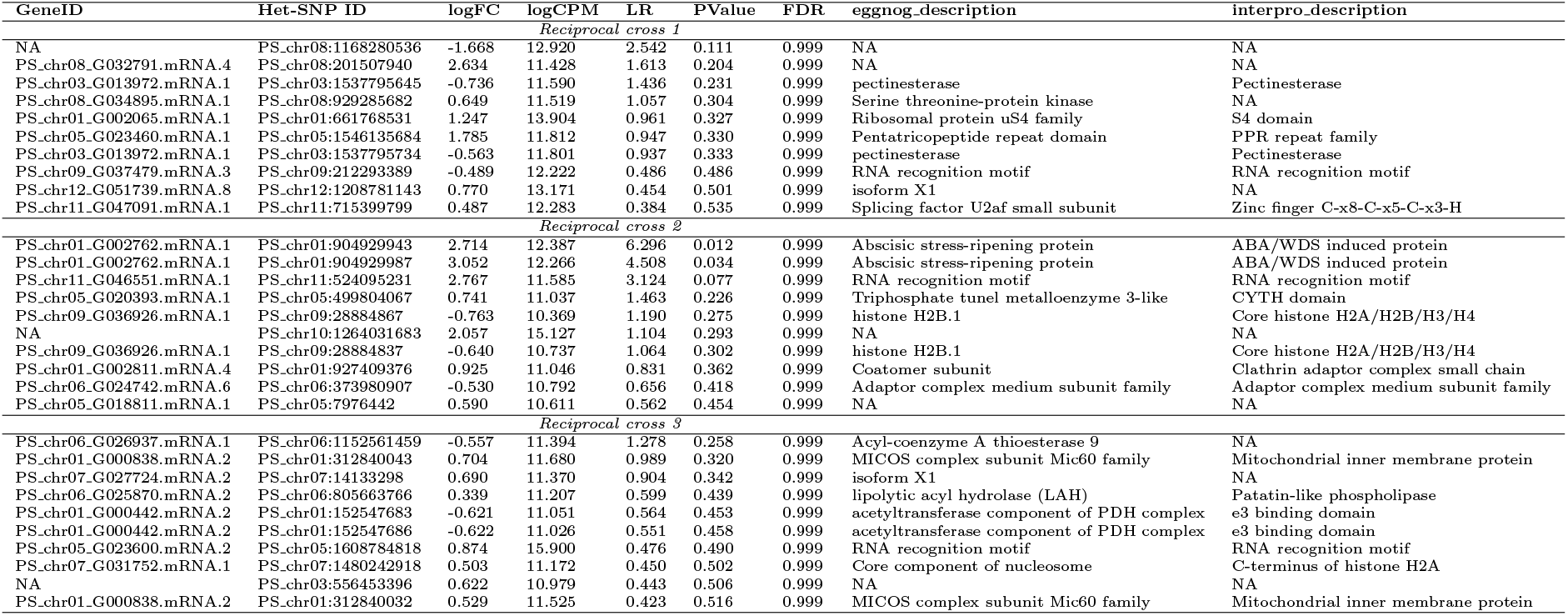
Top edgeR hits. The table lists the ten heterozygous SNPs with the lowest P-values from each reciprocal cross, together with log fold change (logFC), average log counts per million (logCPM), likelihood ratio (LR), P-value, false discovery rate (FDR), and protein annotations from eggNOG and InterPro. Het-SNPs may overlap with multiple mRNAs, but the best matches are shown here.

**Figure 2.**
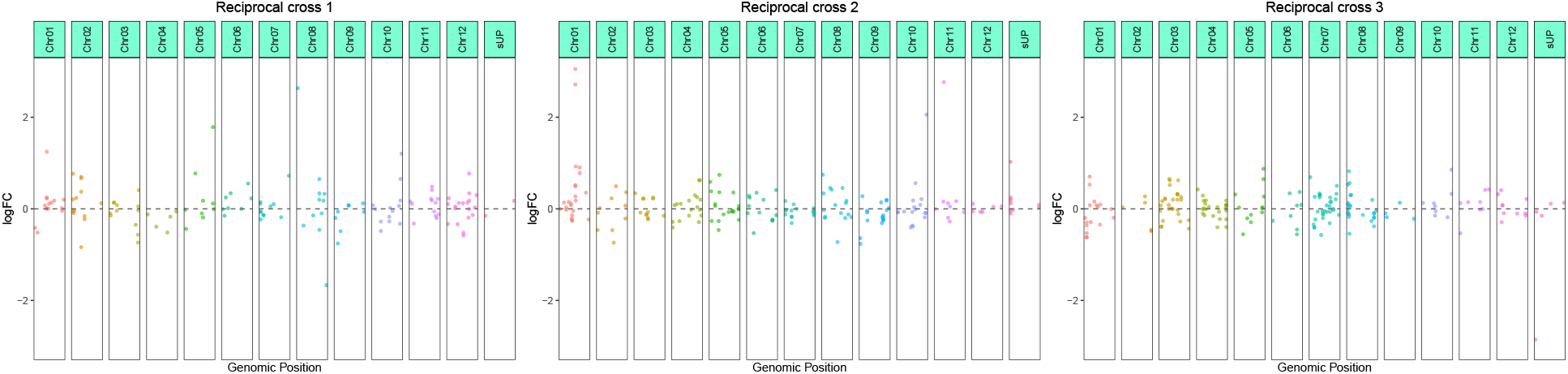
Genome-wide distribution of parent-of-origin expression bias across reciprocal crosses. Manhattan-style plots showing log fold change (logFC) values for all expressed genes across chromosomes in three reciprocal crosses. Each point represents an individual gene positioned according to its genomic coordinate (x-axis) and colored by chromosome. The y-axis shows the logFC between maternal and paternal allele expression, where positive values indicate maternal-biased expression and negative values indicate paternal-biased expression. The dashed horizontal line marks logFC = 0, corresponding to equal parental expression. Chromosomes are displayed sequentially along the x-axis, with the pseudochromosome presented last. This pseudochromosome contains contigs that could not be assigned to any of the established chromosomes.

**Figure 3.**
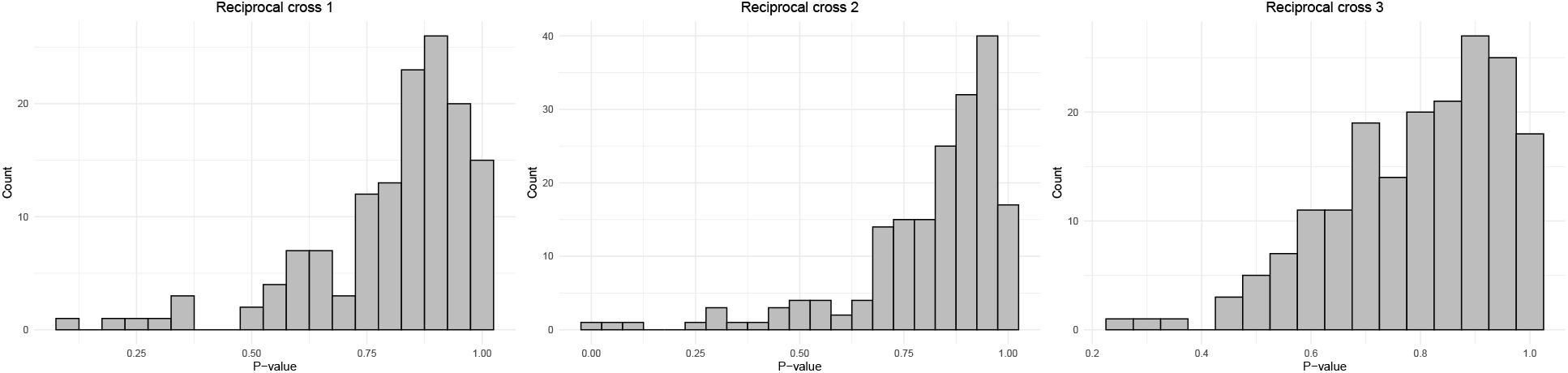
Distribution of raw P-values from genomic imprinting analysis across reciprocal crosses. Histograms show the distribution of raw P-values obtained from edgeR likelihood ratio tests (LRT) for each reciprocal cross (Cross 1, Cross 2, and Cross 3). Each subplot represents one reciprocal cross, with bars indicating the number of genes within a given P-value bin (bin width = 0.05).

**Figure 4.**
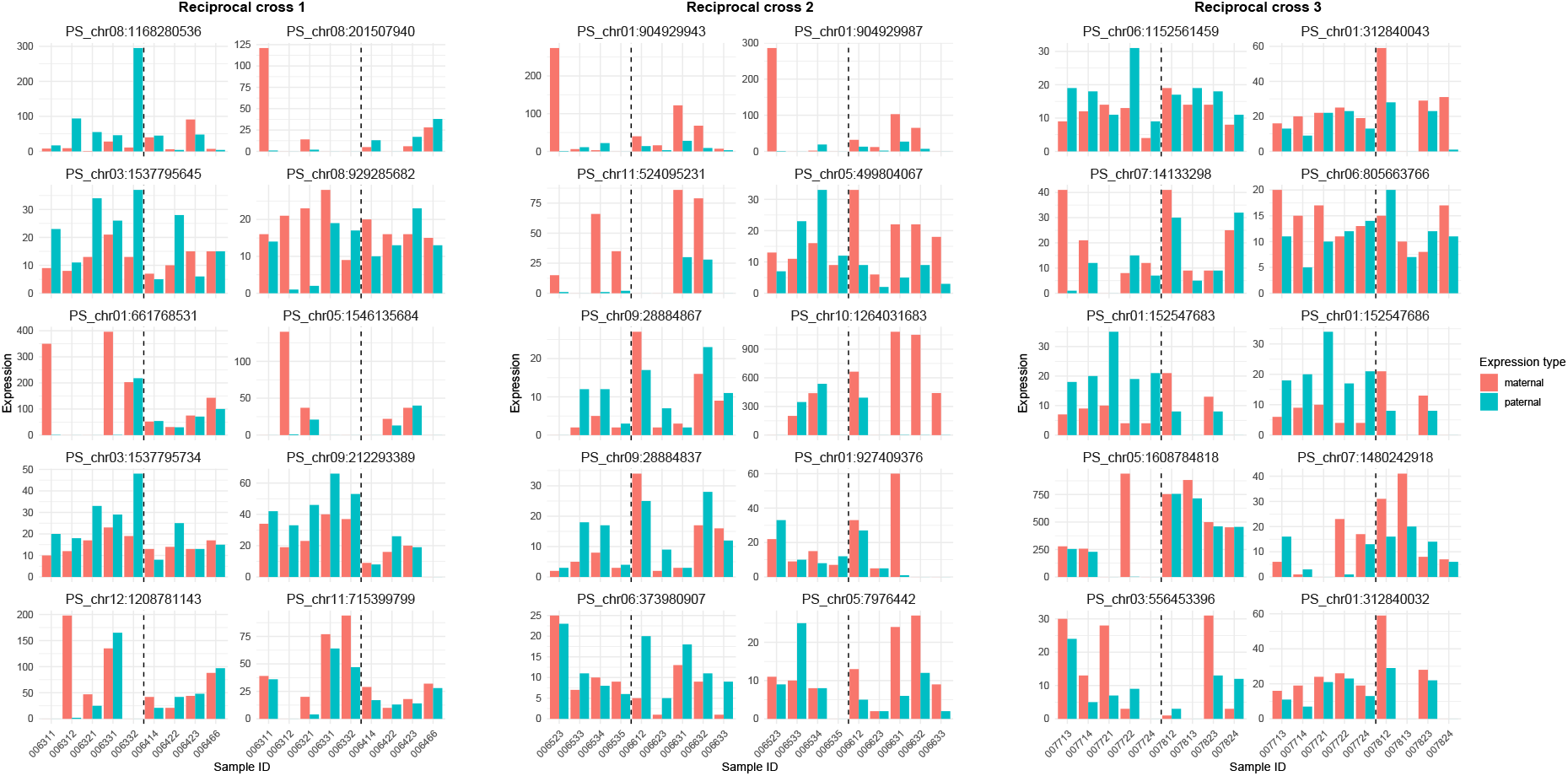
Expression levels for the top 10 het-SNPs with lowest P-values from reciprocal crosses. Bar plots show raw expression values, with each panel corresponding to a single het-SNP. Within each panel, bars represent maternal and paternal expression for individual biological replicates. A black dotted line separates the two reciprocal crosses.

## 4 Discussion

In this study, we aimed to improve understanding on evolutionary origins of genomic imprinting by investigating its potential presence in Scots pine. Evidence of imprinting in Scots pine would further challenge the prevailing view that imprinting arose through convergent evolution, as it would place imprinting in at least three distinct plant lineages: angiosperms, gymnosperms, and liverworts. Such a pattern would instead hint at a shared evolutionary origin for imprinting across land plants, rather than repeated, independent emergence. To investigate this question, we implemented a novel approach for studying imprinting. Our results demonstrate that the use of maternally inherited haploid megagametophyte tissue not only provides a reliable means of identifying parental alleles in the offspring, but also enables the elimination of large numbers of paralogous sites from the data. This approach can be readily adopted in other conifer species and in taxa with similar haploid maternally inherited tissue. We found no strong indications of genomic imprinting from the limited number of heterozygous SNPs available for the analysis in each reciprocal cross, representing less than half a percent of all protein-coding genes in the Scots pine genome. Thus we cannot rule out imprinting for all the genes or in other yet unexplored developmental stages, highlighting the necessity for further research on the subject.

The ability to confidently identify parental alleles and detect genomic imprinting was constrained by several key factors. 1) Limited overlap between the exome capture and RNA-sequencing datasets. Conifer genes are characterized by extremely long introns (Nilsson et al., 2025; Nystedt et al., 2013), whereas exome capture targets relatively short regions spanning a limited number of exons. When combined with the short read lengths of paired-end sequencing data from both exome capture and RNA-sequencing, this substantially reduced the likelihood of overlap between the two datasets. 2) Missing data due to inconsistency and biological variation among replicates. Scots pine seeds, and seeds in general, exhibit variation in size and morphology, leading to differences in library yield and the extent of PCR amplification required to generate sufficient sequencing material. These inconsistencies can result in missing data across replicates, potentially causing true imprinted genes to be missed. Missing data also decreases the overlap between exome capture and RNA-sequencing datasets. 3) Limited heterozygosity. Genomic imprinting is typically investigated using reciprocal crosses between two inbred lines, producing F_1_ offspring with abundant heterozygous sites. In contrast, the Scots pine trees used here were selected from a wild population, since highly homozygous inbred lines are unavailable in species with such long generation times. Conifers also carry a substantial load of lethal and deleterious alleles, which severely limit the success of self-fertilization and render the generation of inbred lines challenging, if not infeasible (Williams and Savolainen, 1996). Furthermore, because many alleles in natural populations segregate at low frequency, most loci are homozygous in any given individual, leading to generally low levels of heterozygosity in Scots pine (Pyhäjärvi et al., 2007). 4) Presence of paralogous regions. The Scots pine genome contains over 49,000 protein-coding genes and more than 155,000 pseudogenes (Nilsson et al., 2025). These numbers are largely explained by local gene duplications, which are a major source of genetic novelty in conifers (Nilsson et al., 2025). However, the abundance of duplicate genes complicates genetic analyses in Scots pine. Reads mapping to paralogous regions can introduce substantial biases in various analyses, including studies of genomic imprinting. To mitigate these issues, we applied very strict genotype filtering step, as we could not reliably distinguish heterozygous maternal genotypes from paralogous sequences. We also removed multi-mapped reads from the RNA-sequencing data, but many of the heterozygous SNPs we identified may still be false positives, arising from incorrect mapping of paralogs. The two het-SNPs with low P-values showing inconstant expression patterns for genomic imprinting may have arisen from this mapping ambiguity.

These limiting key factors can be addressed in future experiments by incorporating the following changes: Exome capture efficiency can be improved by employing short-read sequencing platforms capable of generating longer reads (e.g., 2×300 bp or 2×500 bp) and by increasing the amount of species-specific c0t-1 DNA used during hybridization. The use of species-specific c0t-1 DNA has been observed to reduce the capture of various repetitive DNA sequences, thereby increasing the capture of unique exon sequences compared to the use of commercial universal blocking agents (Kesälahti et al., 2025). Another strategy is to use whole-genome sequencing; however, this approach is not viable for species with large, highly repetitive genomes without major workarounds. Pairing long-read sequencing (PacBio, Nanopore) or extended short-read approaches with Iso-Seq or Nanopore Direct RNA Sequencing enables full-length transcript capture and more accurate RNA-seq mapping, while the longer DNA reads from PacBio and Nanopore also improve genomic alignment. Together with increased sequencing depth, these improvements can substantially boost dataset overlap and thereby strengthen parental-allele identification. The use of sequencing methods with longer reads can also help address the paralog issue, as longer reads are more likely to map accurately to their corresponding sequences in the reference genome. Another strategy to mitigate issues caused by paralogs is to identify and remove highly paralogous clusters from the reference. However, this approach must be applied carefully, as it could lead to exclusion of imprinted genes from the dataset. Data gaps can be mitigated by increasing the number of replicates and discarding seeds with low DNA or RNA extraction yields, thereby generating more evenly sized libraries. The lack of heterozygosity can be addressed in some species by performing reciprocal crosses between genetically or geographically distant populations. Increased heterozygosity also has the potential to mitigate the paralog issue, as a higher number of SNPs would be present between alleles derived from different populations. To definitively confirm the imprinting status of a gene, methylation data is essential. Epigenetic modifications that regulate genomic imprinting cannot be detected through standard sequencing data alone, as they involve chemical changes to the DNA that influence gene expression without altering the underlying genetic sequence. Majority of these improvements were out of scope for this small scale pilot experiment. However, establishing whether genomic imprinting occurs in conifers will require further investigation, incorporating the improvements mentioned above, to clarify its evolutionary origin and potential role in conifer adaptation.

## 5 Conclusions

We investigated whether genomic imprinting, an epigenetic process that causes parent-of-origin–specific gene expression, occurs in Scots pine and thus possibly in other conifer species. Our goal was to gain insight into the evolutionary origins of imprinting and to provide evidence on concerning whether it evolved independently in angiosperms and animals through convergent evolution or reflects a common ancestral origin. We found no direct evidence of genomic imprinting, given the limited number of heterozygous SNPs available for analysis. Consequently, the data did not allow us to draw definitive conclusions about the presence of imprinting in conifers. Nonetheless, we show that genomic imprinting can be examined in conifers by utilizing maternally derived haploid megagametophyte tissue to identify parental alleles and to eliminate large quantities of paralogous sites from the data. Our study faced several critical limitations, with the prevalence of paralogous regions due to gene duplication and the minimal overlap between exome capture and RNA sequencing datasets being the most significant. We suggest several improvements to the study design to overcome these challenges, as we consider it important to confirm or refute the existence of genomic imprinting in conifers. Establishing this will enhance our understanding of its evolutionary history and evaluate its possible contribution to the adaptability of conifer species.

## 7 Data availability statement

Raw RNA-sequencing and exome capture reads have been deposited in the NCBI Sequence Read Archive under BioProject accession PRJNA1440209. During the peer-review process, these data are accessible via the following reviewer link: https://dataview.ncbi.nlm.nih.gov/object/PRJNA1440209?reviewer=eeli4rc5o8j720hhve00m5efuq. All bioinformatics scripts and processed allelic count tables used in the edgeR analysis, including a modified version of the pipeline developed by Wyder et al. (2019), are available via the Figshare reviewer portal: https://figshare.com/s/0d5f6bdcce89fd2fd8f4. All accession numbers and repository links will be made publicly available upon publication.

## 8 Author contributions

Tanja Pyhäjärvi conceived the study and designed the experiments. Sandra Cervantes and Robert Kesälahti performed the wet laboratory work. Robert Kesälahti designed the analytical approach and performed the data analyses with help from S. Cervantes, Alina K. Niskanen and T. Pyhäjärvi. R. Kesälahti wrote the manuscript with help and comments from S. Cervantes, A. Niskanen and T. Pyhäjärvi.

## 9 Acknowledgments

We thank Komlan Avia for designing the reciprocal crosses and the Natural Resources Institute Finland for carrying them out. We also thank Soile Alatalo for her expertise and assistance with the wet laboratory work. The authors acknowledge CSC—IT Center for Science, Finland, for providing computational resources.

## 9.1 Study funding

This work was supported by the following grants: Research Council of Finland (293819 and 319313 to Tanja Pyhäjärvi), Emil Aaltonen Foundation (Nuoren tutkijan apuraha to Robert Kesälahti) and Finnish Cultural Foundation (Working grant to Robert Kesälahti).

